# Fetal gut colonization: meconium does not have a detectable microbiota before birth

**DOI:** 10.1101/2021.02.17.431710

**Authors:** Katherine M. Kennedy, Max J. Gerlach, Thomas Adam, Markus M. Heimesaat, Laura Rossi, Michael G. Surette, Deborah M. Sloboda, Thorsten Braun

**Author notes:** shared senior authorship (Deborah M. Sloboda, Thorsten Braun). Corresponding Authors, D. M. Sloboda McMaster University, Department of Biochemistry and Biomedical Sciences 1280 Main St West, Hamilton, Canada, T. Braun, Charité – University Medicine Berlin, Departments of Obstetrics and Division of Experimental Obstetrics Charité Campus Virchow, Berlin, Germany.

## Abstract

Microbial colonization of the human intestine impacts host metabolism and immunity, however when colonization occurs is unclear. Although numerous studies have reported bacterial DNA in first-pass meconium samples, these samples are collected hours to days after birth. We investigated whether bacteria could be detected in meconium prior to birth. Fetal meconium (n = 20) was collected by rectal swab during elective breech Cesarean sections without labour prior to antibiotics and compared to technical and procedural controls (n = 5), first-pass meconium (neonatal meconium; n = 14), and infant stool (n = 25). Unlike first-pass meconium, no microbial signal distinct from negative controls was detected in fetal meconium by 16S rRNA gene sequencing. Additionally, positive aerobic (n = 10 of 20) and anaerobic (n = 12 of 20) clinical cultures of fetal meconium (13 of 20 samples positive in at least one culture) were identified as likely skin contaminants, most frequently *Staphylococcus epidermidis*, and not detected by sequencing in most samples (same genera detected by culture and sequencing in 2 of 13 samples with positive culture). We conclude that fetal gut colonization does not occur before birth, and that microbial profiles of neonatal meconium reflect populations acquired during and after birth.

## Introduction

Microbial colonization of the human intestine is a key developmental process as the order and timing of microbial exposures shape the development of the gut microbiome^1^ and impact host metabolism and immunity later in life.^2^ In humans, maturation of intestinal barrier function and immunity both occur prenatally.^3^ The fetal intestine is more permeable to macromolecules^4^ and less tolerant of antigens^5^ than that of term infants. Transfer of maternal IgG across the placenta and uptake in the fetal intestine increase near term gestation,^6^ shaping neonatal gut immune responses after birth.^7^ Thus the intrauterine environment has the capacity to shape health well beyond fetal life, and can influence long term health trajectories,^8^ and recently it has been suggested that early-life colonization with specific microbes can predict health outcomes including asthma^9^ and obesity.^10^ To understand the mechanisms by which microbial colonization influences health later in life we must know when colonization occurs. Several groups using sequencing-based methods have reported bacterial DNA in the placenta^11,12^ and amniotic fluid^13^ and have suggested that this reflects microbial populations that initiate gut colonization *in utero*.^14^ However, recent studies accounting for the high risk of contamination in low-biomass samples^15^ have failed to detect a placental^16–18^ or amniotic fluid microbiome.^19–21^ Thus, this issue remains highly controversial.

As neonatal (“first-pass”) meconium is formed prior to birth, it has been used as a proxy for the *in utero* environment,^22,23^ but this does not account for microbial acquisition that occurs during and/or immediately after birth. Recent metagenomic evidence^24^ and previous culture data^25^ show a correlation between the time from birth to collection and bacterial detection in neonatal meconium. Only one previous study has evaluated the presence of microbes in fetal meconium prior to birth,^26^ and found that at mid-gestation the majority of fetal meconium bacterial profiles did not differ from procedural and kidney controls.^26^ Those that did differ were dominated by *Micrococcus* and *Lactobacillus* species, likely originating from the maternal cervicovaginal microbiota during sample collection.^27^ Here, we characterized the bacterial profiles of human fetal meconium prior to birth. We show that unlike neonatal meconium, fetal meconium is indistinguishable from negative controls, indicating that colonization of the human gut likely does not occur prior to birth. These data significantly extend our understanding of the establishment of our intestinal microbiome and shed light on which early life influencers may impact postnatal gut health.

## Results

### Participant characteristics

To investigate the possible colonization of the fetal gastrointestinal tract *in utero*, we analyzed meconium samples collected from 20 term fetuses during caesarean section prior to birth after an intensive method establishment (Table 1). Fetal meconium was sampled by rectal swabbing during elective caesarean section deliveries with no signs of labour, preterm labour, or rupture of membranes to prevent vertical transmission during labour (Fig. 1; Supplementary Fig. 1). As others have previously detected a microbiome in first-pass (neonatal) meconium,^2,22^ we also included neonatal meconium from term deliveries and infant stool samples collected at 6 months of age as positive controls. We included multiple negative controls: a swab exposed to operating room air during caesarean delivery (sampling negative – collected in triplicate for sequencing and cultures during collection of M213), genomic prep reagents either exposed to PCR hood air during sample preparation or not exposed (extraction negatives), and V3-V4 PCR amplifications without added template DNA (PCR negative).

**Table 1.** Pa11icipant char·acteristics.

|  | Fetal<br>(n = 20) | Neo<br>(n = 14) |
| --- | --- | --- |
| Maternal age, years<br>mean (SD) | 32.3 (5.1) | 33.2 (4.4) |
| Gestational age, days *<br>mean (SD) | 271.4 (5.6) | 276.7 (6.0) |
| Birth weight, grams<br>mean (SD) | 3240 (380) | 3384 (557) |
| Sex |  |  |
| Female | 14 (70) | 7 (47) |
| Male | 6 (30) | 8 (53) |
| Delivery, n (%) <sup>+</sup> |  |  |
| Vaginal | 0 (0) | 9 (60) |
| Caesarean | 20 (100) | 8 (40) |

p < 0.05 indicated by \* (t-test) or <sup>+</sup> Chi-squared

**Figure 1.**
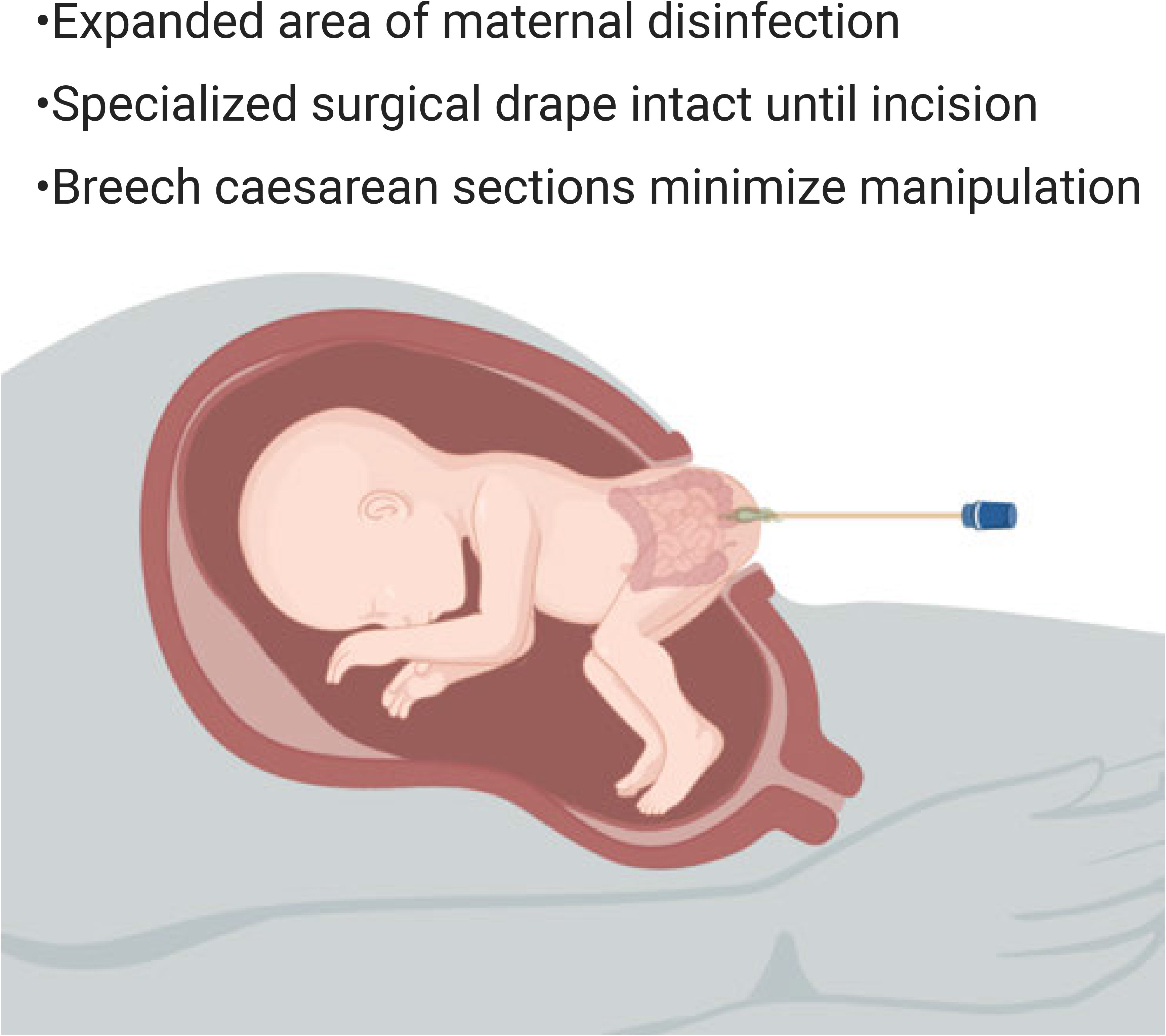
Diagram of collection method for fetal meconium samples. Following advanced physical disinfection of maternal skin from maternal armpits to knees, the surgical area was covered using the specialized sterile drape, with film remaining intact until the incision. After caesarean section and exposure of the fetal buttocks prior to birth, meconium was rectally sampled using sterile eSwabs™.

Fetal and neonatal participants did not differ significantly in maternal age or birth weight. The gestational age of fetal participants at birth was less than that of neonatal participants (p=0.0010; Welch’s t-test), likely due to differences in mode of delivery (p=0.00029, **χ**-squared).

### Clinical culture results

Fetal meconium samples and the sampling negative control were cultured under both aerobic and anaerobic conditions (Table 2). Of the 20 fetal meconium samples, 7 were negative for both aerobic and anaerobic cultures after 120 hours, as was the sampling negative control. Additionally, 3 samples had negative aerobic cultures but positive anaerobic cultures (M208, *Staphylococcus epidermidis*; M210, *Propionibacterium acnes*; M217, *Propionibacterium acnes*) and 1 sample had a negative anaerobic culture but positive aerobic culture (M219, *Staphylococcus epidermidis*). Despite growth of *Staphylococcus epidermidis* under both aerobic and anaerobic conditions only 3 meconium samples had positive cultures under both conditions, while 5 had positive cultures under only one. As *Propionibacterium acnes* and coagulase-negative Staphylococci are common skin contaminants in clinical cultures, we attempted to confirm positive results in sequencing data.

**Table 2.** Fetal meconium culture results.

| ID | Anaerobic culture |  | Aerobic culture |  |
| --- | --- | --- | --- | --- |
|  | Result | Incubation<br>Time (hrs) | Result | Incubation<br>Time (hrs) |
| M201† | Staph. epidermidis | 18.64 | Staph. epidermidis | 23.2 |
| M202 | Staph. epidermidis | 18.68 | Staph. lugdunensis | 20.67 |
| M203* | Staph. epidermidis | 20.12 | Staph. epidermidis | 14.92 |
| M204*† | Staph. epidermidis | 18.59 | Staph. epidermidis | 17.09 |
| M205† | negative | 120.07 | negative | 120.07 |
| M206 | Staph. saprophyticus | 26.09 | Staph. saprophyticus | 17.96 |
| M207† | Staph. epidermidis | 19.06 | Staph. lugdunensis | 18.69 |
| M208 | Staph. epidermidis | 22.05 | negative | 120.08 |
| M209 | negative | 120.07 | negative | 120.08 |
| M210 | Propionibacterium acnes | 79.21 | negative | 120.08 |
| M211 | negative | 120.08 | negative | 120.09 |
| M212 | Staph. epidermidis | 19.94 | Staph. hominis | 24.29 |
| M213 | Propionibacterium acnes | 102.77 | Staph. capitis | 36.68 |
| Neg | negative | 102.02 | negative | 120.03 |
| M215 | negative | 120.13 | negative | 120.13 |
| M216 | negative | 120.16 | negative | 120.15 |
| M217 | Propionibacterium acnes | 66.24 | negative | 120.17 |
| M219 | negative | 120.08 | Staph. epidermidis | 17.41 |
| M221 | negative | 120.08 | negative | 120.08 |
| M222 | negative | 120.08 | negative | 120.09 |
| M223 | Propionibacterium avidum | 80.85 | Propionibacterium avidum | 99.05 |

\* monochorionic-diamniotic twins; † not breech presentation

### 16S rRNA marker gene sequencing

We assessed bacterial DNA present in samples using 16S rRNA gene sequencing of combined variable 3 and 4 region (V3V4) amplicons from 30 cycles of PCR amplification. All meconium samples and negative controls were additionally sequenced after 40 PCR cycles to confirm the presence of any genera detected at 30 cycles. We first looked at the number of amplicons sequenced in each sample (read count). Fetal meconium samples were dominated by host-associated reads before removal of host-associated taxa (pruning) (median read count of 109.5; min = 23; max = 1316; n=20), after which they had a median read count of 76.5 (min = 16, max = 202, n=20). Read counts of neonatal meconium samples were much more variable: two samples did not have any reads, and the remaining samples had a median read count of 130 (min = 4, max = 63079, n=12) before pruning and a median read count of 191 after (min = 9, max = 63079, n=11). Negative controls had a median of 74 reads (min = 6, max = 396) before pruning and a median of 46 reads after (min = 6, max = 99). As expected, infant stool samples had much higher read counts than other samples (median read count of 53207; min = 5505; max = 105212; n = 25) before and after pruning.

Looking at within-sample diversity, fetal meconium had lower alpha diversity (Observed ASVs, Shannon index, Simpson’s index, Fig. 2A) than infant stool (p = 0.0016, p = 0.0001, and p = 0.0086 respectively). Neonatal meconium alpha diversity did not differ from that of infant stool (p = 0.7503, p = 0.6401, and p = 0.7762 respectively). To investigate differences in overall community composition between samples (beta-diversity) we performed principal coordinate analysis (PCoA) of Bray-Curtis dissimilarity and found fetal meconium samples cluster with negative controls, while neonatal meconium samples were more variable and more similar to infant stool samples (Fig. 2B). Overall beta-diversity of fetal meconium was indistinguishable from that of negative controls (Fig. 2C) as Bray-Curtis dissimilarity between samples was similar within negative controls, within fetal meconium, and between negative controls and fetal meconium. Compared to fetal meconium, neonatal meconium was more dissimilar to negative controls (p<0.001).

**Figure 2.**
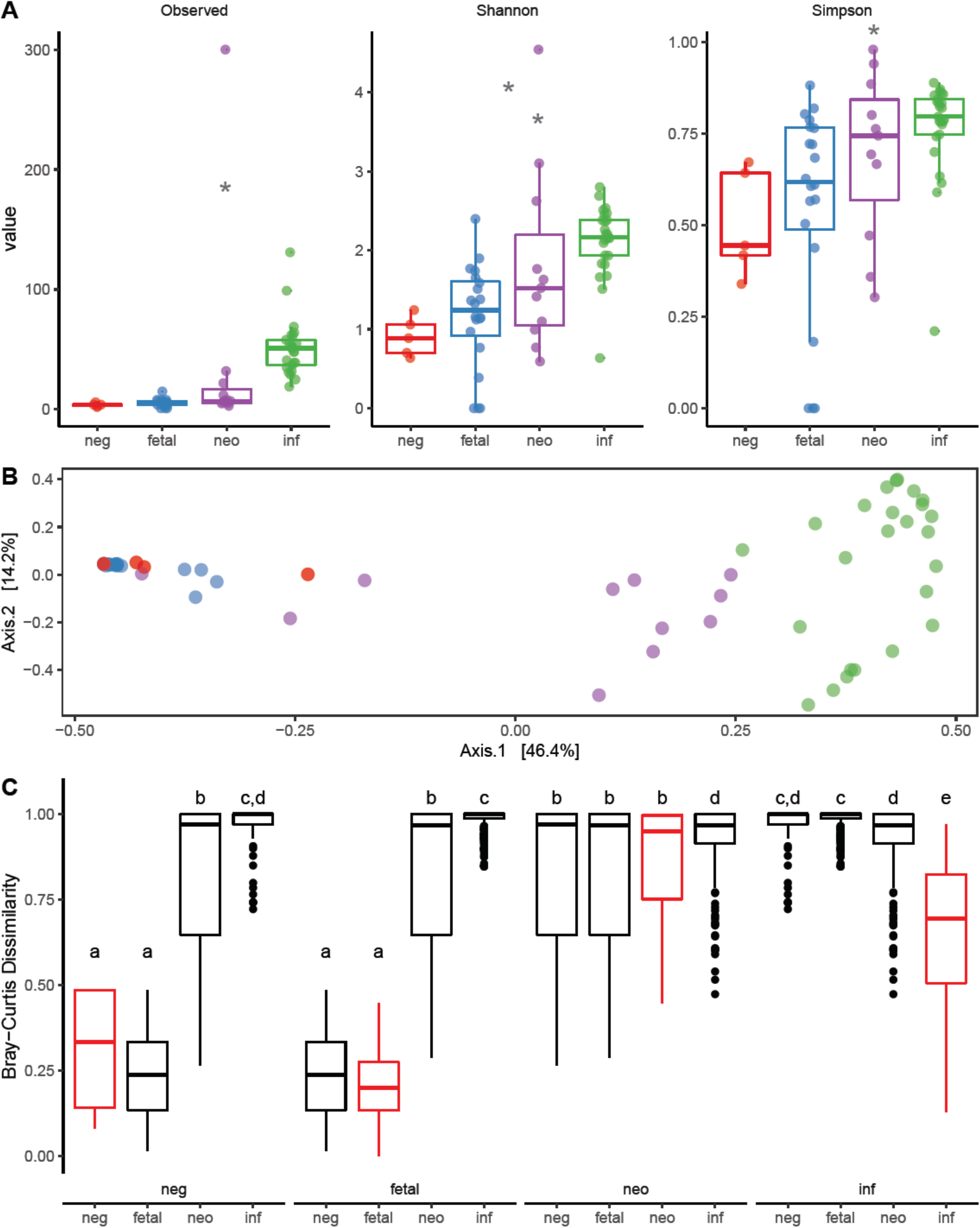
Fetal meconium alpha and beta diversity do not differ from those of sampling negative controls. a, Boxplots of alpha diversity measures by sample type (negative control (neg, n = 5), fetal meconium (fetal, n = 20), neonatal meconium (neo, n = 14), and infant stool (inf, n = 25). Significance assessed by a linear mixed model, with sample type as a fixed effect and participant ID as a random effect (p < 0.05 indicated by *). b, Principal coordinate analysis (PCoA) of Bray-Curtis distances at genus level shows fetal meconium samples (blue dots) cluster with sampling negative controls (red), indicating their community composition is similar, while neonatal meconium samples (purple) are more variable. c, Boxplots of Bray-Curtis dissimilarities within (red) and between (black) sample types shows that fetal meconium samples less dissimilar to each other and similar to negative controls. Significance assessed by linear mixed model with sample type comparison as a fixed effect (i.e. neg-neg, neg-fetal, etc.) and participant ID as a random effect. Boxplot center line, median; box limits, upper and lower quartiles; whiskers, 1.5x interquartile range; points, outliers.

The most prevalent genera detected in fetal meconium samples were *Halomonas* (20/20 samples), *Rhodanobacter* (19/20, only detected in 1 of 2 sequencing runs for 4 samples), and *Pseudomonas* (15/20, only detected in 1 of 2 sequencing runs for 6 samples), all of which were also detected in negative controls (Fig. 3). The only genera detected in more than one fetal meconium sample and not detected in negative controls were *Bacteroides* (4/20 samples, only detected in 1 of 2 sequencing runs for 2 samples) and *Staphylococcus* (4/20 samples, only detected in 1 of 2 sequencing runs for 2 samples). *Bacteroides* was only consistently detected across sequencing runs and with both 30 and 40 cycles of amplification in 1 of 20 samples (M202; Supplementary Fig. 2). *Staphylococcus* was not consistently detected with both 30 and 40 cycles of PCR amplification in any sample and was only detected by sequencing in 2 of 11 samples with positive *Staphylococcus* culture results (M201 positive for *S. epidermidis* and M207 for both *S. epidermidis* and *S. lugdunensis*). The only other genus consistently detected in a sample with both 30 and 40 cycles of amplification and across sequencing runs was *Escherichia*/*Shigella* (M202, M203, M207) which was also detected in an extraction negative control (Supplementary Fig. 2). Despite culture results positive for *Propionibacterium* species, no members of this genus were detected in sequencing data from any fetal meconium sample.

**Figure 3.**
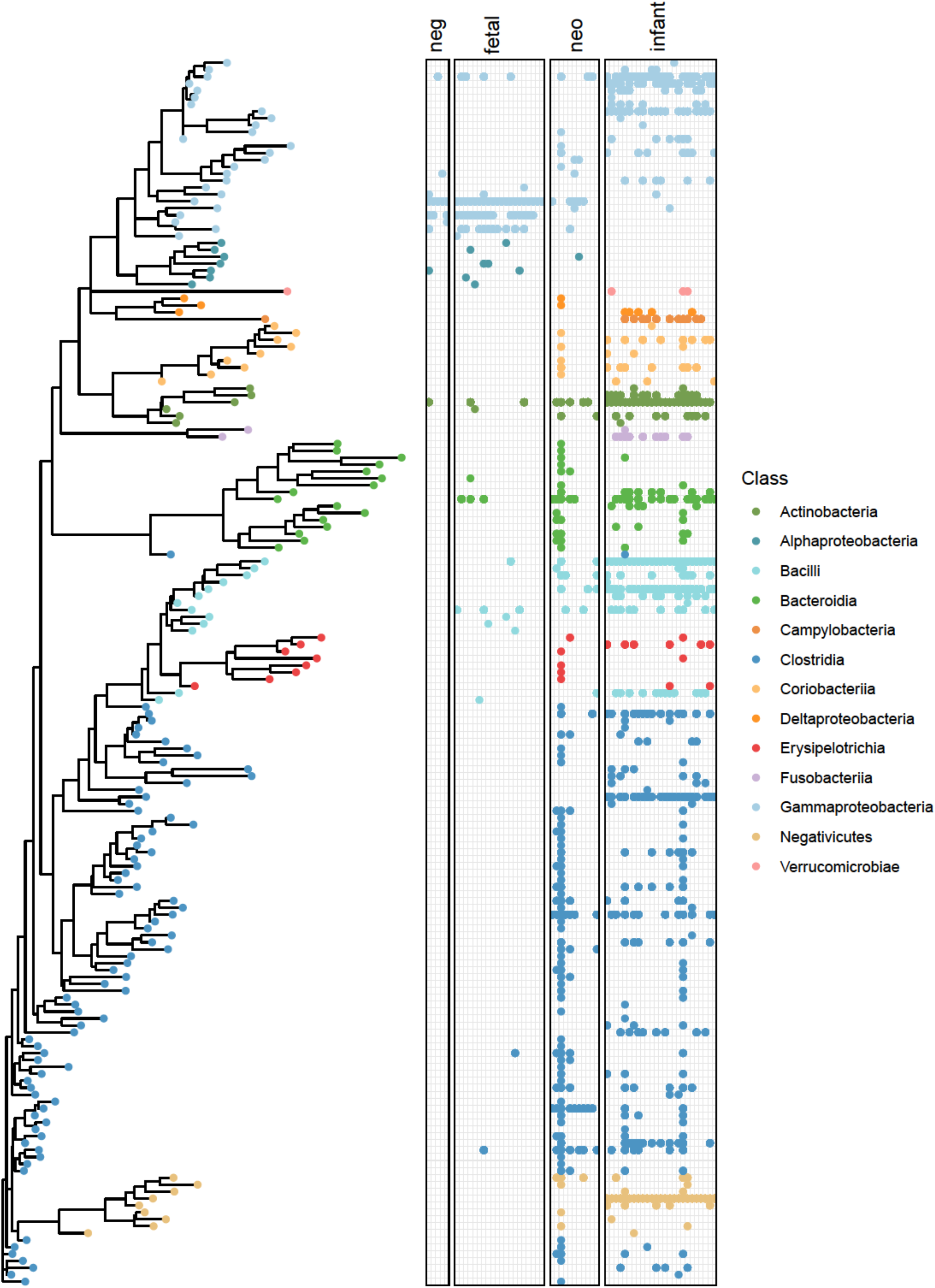
Neighbour-joining phylogenetic tree of all genera and presence/absence in each sample. Taxonomy was assigned using the RDP Classifier against the Silva 132 reference database. Dots indicate presence of the genera at any abundance in the given sample. Gammaproteobacteria prevalent in fetal meconium (fetal) are also prevalent in sampling and extraction negative controls (neg). Microbial profiles of first-pass neonatal meconium (neo) share more genera with those of infant stool (inf) than fetal meconium.

## Discussion

The role of the microbiome in controlling host metabolism and its relationship to metabolic dysfunction and obesity^28^ has led investigators to question whether our microbial signatures early in life could be used to predict chronic disease risk.^29^ Because of this, how early in life host-microbe interactions are established has become a topic of intense investigation. In neonates, several microbial species are known to regulate intestinal function^30^ and are key in immune development.^31^ Recent studies reporting colonization may be initiated *in utero*^11,12,14^ have been the subject of vigorous debate and have been criticized for potential contamination.^16,17,21^ Despite our extensive sample collection optimization to reduce potential contamination (see Methods and Supplementary Fig. 1) more than half of fetal meconium samples had at least one culture test positive for a likely skin contaminant (22/40 total cultures), highlighting the difficulty in avoiding contamination in these types of studies. Contaminating bacteria or bacterial DNA may be introduced by maternal skin or blood during sampling, the environment or investigators during sample handling and culturing, or reagents during DNA extraction and PCR. Negative controls are therefore necessary at each of these stages to rule out contamination. We analyzed fetal meconium collected immediately prior to birth, and compared term fetal meconium to appropriate negative controls,^32^ to neonatal meconium and to infant stool samples. This was a particular strength of our study. Additionally, we used a combination of clinical culture and 16S rRNA gene sequencing to identify a fetal microbial signature. Despite all efforts, we were unable to detect a microbial signature in fetal meconium that was distinct from negative controls. Overall, the lack of a consistent bacterial signal in our data indicates that fetal meconium does not have a microbiome prior to birth.

Staphylococci were the dominant genus in culture results (17 cultures) and *S. epidermidis* was the most prevalent species (11 cultures). In our pilot cohort postoperative swabs of maternal skin around the incision were also positive for *S. epidermidis*. While it is not possible to know if positive cultures were the result of contamination by skin microbes, the absence of cultured genera in sequencing data make this a likely possibility. Although *S. epidermidis* DNA has also been reported in amniotic fluid^14^ and neonatal meconium, ^24^ it is most frequently associated with the skin microbiota^33^ suggesting that despite especially stringent efforts to control for contamination some element of contamination may always exist with current technologies and protocols. This makes the strong case for ensuring that a robust set of controls are in place when performing these types of investigations.

Although sequencing cannot prove the absence of bacterial DNA in a sample, we used robust negative controls and technical replicates to distinguish contamination signals from stochastic sequencing noise. While previous studies have reported the existence of bacterial DNA in amniotic fluid and the placenta, these studies lacked sequencing data from negative controls and therefore cannot rule out contamination.^15^ In our study, we included negative controls collected during sampling, extraction, and PCR amplification. Inclusion of these negative controls allowed us to identify the most prevalent and abundant genera as likely contaminants introduced during extraction (*Halomonas* and *Rhodanobacter*) or sampling (*Pseudomonas*). Additionally, we included neonatal meconium as a positive control of our extraction and amplification of bacterial DNA from these low-biomass samples.

Even when sequencing data from appropriate negative controls are included, low-biomass samples are especially sensitive to stochastic amplification^34^ and sequencing^32^ noise. Determination of a true bacterial signal requires its presence across technical replicates. In this study, we ran the PCR products of each amplification on two separate sequencing runs. Excluding likely contaminants, almost all genera detected were only found in one of two sequencing runs for each sample. *Bacteroides* was the only genus consistently detected across technical replicates, and this was only true in 1 of 20 fetal meconium samples. Thus, when comparing across sequencing runs, we found that the bacterial signals in fetal meconium that are not also detected in negative controls are likely due to stochastic sequencing noise.

In terms of overall community composition, fetal meconium beta-diversity was similar to that of negative controls and lower than that of neonatal meconium. Neonatal meconium community composition was highly variable with some samples more closely resembling infant stool. Thus, microbial profiles of neonatal meconium reflect populations acquired during and after birth and not populations that exist prior to birth.

To our knowledge, this is the first study to investigate the meconium microbiome in term human neonates prior to birth and this is also the first study to control for both contamination during sampling and DNA extraction and stochastic noise during sequencing. While it is not possible to prove the absence of bacteria in fetal meconium prior to birth, our data do not support *in utero* colonization. These data suggest that colonization more likely occurs either during birth via maternal skin/vaginal/fecal seeding, or post birth via environmental seeding.

## Methods

### Study design and sample collection

#### Implementation phase

A pilot cohort of 22 participants was recruited to optimize sample collection methods to minimize risk of contamination and comprised two pilot cohorts as described below (Pilot 1a and 1b). Only caesarean deliveries were included to avoid the vertical transmission of bacteria during a vaginal birth, during which the child is exposed to the maternal bacterial flora. In addition to the duration of the birth process itself, a further disadvantage of spontaneous delivery is the variable time within which the newborn engages its first bowel movement (neonatal meconium - which is of prenatal origin). Study inclusion was further restricted to elective caesarean sections prior to labour due to potential microbial influences on unplanned section deliveries and to guarantee the presence of an operator familiar with the study protocol and an equally trained research personnel who was able to carry out the immediate transport and further processing of the sample in the nearby laboratory. Breech presentations were preferred to ensure that the rectum was immediately accessible for sampling after sectioning and before delivery and to minimize manipulation of the child before sampling (Fig. 1). This ensured a reduced risk of contamination as a swab could be taken before the neonate was fully removed from the uterus. Finally, to avoid false-negative results the preoperative prophylactic antibiotic was administered after meconium sample collection.

#### Pilot cohort 1a (establishment of sampling procedures)

Swabs with nylon flock fiber (eSwab ™, Copan Diagnostics Inc.) were used to collect fetal meconium samples. In the event of a meconium leakage (spontaneous as a side effect of breech presentation), a sample was either taken up with a swab, or a careful rectal smear was performed. Pilot samples were transferred to blood culture bottles (BD BACTEC™) and brought to the Charité University Berlin laboratory for culturing. The inoculation of the blood culture bottles was initially not carried out under sterile conditions, but the use of a sterile workbench was quickly considered and tested (from #4 onward, not continuously). An amniotic fluid sample (#9) and fetal perianal swabs (#6, #7) were also cultured to identify possible sources of contamination. The Head of Clinical Microbiology from Labor Berlin was consulted for an external review of the study design and protocol. Of the 16 pilot meconium cultures (8 aerobic + 8 anaerobic), 11 showed bacterial detection (68.8%), 2 of which were more than one species (#2, #10). Only 1 of the 8 pilot samples (#5) had both anaerobic and aerobic negative cultures (12.5%), 3 cases (#2, #6, #8) had only anaerobic positive cultures (37.5%) and 4 cases (#3. #4, #7, #10) had both anaerobic and aerobic positive cultures (50%). Positive cultures had a median incubation time of 20 hours until detection. The 13 detected microbial populations could be assigned to the following species in decreasing frequency: *Staphylococcus epidermidis* (5), *Staphylococcus lugdunensis* (3), *Staphylococcus capitis* (2), *Staphylococcus caprae, Propionibacterium acnes*, *Propionibacterium avidum* (1 each). The amniotic fluid sample showed no contamination. *Staphylococcus epidermidis* (both anaerobic and aerobic, #7), *Enterobacter aerogenes* (anaerobic, #6), and *Staphylococcus lugdunensis* (aerobic, #6) were detected in perianal swabs. Maternal skin was considered as the potential source of contamination, and therefore a second pilot study was performed to minimize skin contamination during sampling.

#### Pilot cohort 1b (#11 - #22)

To investigate maternal skin as a potential source of contamination, we cultured a set of pilot swabs of maternal skin around the caesarean section area both preoperatively after disinfection and postoperatively after the end of wound care (#11). These samples were collected and cultured as above in pilot 1a. Anaerobic preoperative swab cultures were positive for *Staphylococcus lugdenensis*, *Bacillus licheniformis*, and *Clostridium perfringens*. Postoperative swab cultures were positive for *Staphylococcus epidermidis* (both anaerobic and aerobic) and *Bacillus licheniformis* (anaerobic) and *Staphylococcus lugdunensis* (aerobic). To reduce contamination from maternal skin microbes, we expanded the area of disinfection with Softasept^R^ N coloured from the maternal armpits to her knees. To reduce the chance of contamination due to amniotic fluid dampening the surgical draping, a new model of draping was designed and used (from #15 on) that had a wider design, better adhesive properties, and a thinner film. The thinner film helped in cases where the mother had repeated caesarean sections, where the prior model sometimes required the film to be slit to visualize the existing scar. The film of the new perforated sheet remained intact and the aseptic area remained undisturbed up to the skin incision. Lateral drapes were also introduced (from #16 on) and the remaining pilot samples were used to observe the resistance of the seal, to identify possible points susceptible to film detachments (#17 - #22).

#### Principal Study cohort

Participants (n=19 mothers) were recruited during routine antenatal counselling when patients presented with planned caesarean section and breech presentation at 36+0 weeks gestation. The study protocol was reviewed and approved by Charité ethics committee (EA 04/059/16). Informed consent was obtained from all study participants.

Advanced physical disinfection of maternal skin was performed (as described in pilot 1b), covering the area from maternal armpits to knees and the surgical area was covered using the specialized sterile drape, with film remaining intact until the incision. After caesarean section and exposure of the fetal buttocks prior to birth, meconium was rectally sampled using sterile eSwabs™. At least one operating room negative control (swab exposed to operating room air) was also taken. Triplicate samples were collected, of which two were immediately cultured (anaerobic and aerobic) and one was flash frozen in liquid nitrogen for future sequencing.

#### Neonatal meconium and infant stool sample collection

Neonatal meconium and infant stool samples used as positive controls were collected as part of a separate study (unpublished). Neonatal meconium samples were collected from diapers within 48 hours of birth and frozen at −80℃ prior to processing for sequencing. Infant stool samples were collected at 6 months of age by parents and frozen at −20℃ before being stored at −80℃ prior to processing for sequencing. Neonatal meconium and infant stool samples were not included in bacterial culturing.

### Bacterial culturing

Samples were transferred in a sterile hood to anaerobic and aerobic point bottles (BD Bactec system™) and supplemented (BD BACTEC ™ FOS ™) to improve the growth conditions for more demanding bacterial species. Cultures were maintained for a maximum of 120 hours. Positive cultures were sub-cultured on agar plates and identified using MALDI-TOF (VitekMS, bioMérieux).

### DNA extraction and amplification

Sample aliquots were transferred to genomic prep tubes in a sterile PCR hood using sterile biopsy punches and gloves were changed between handling each sample. In addition to the sampling negative control collected in the operating room, an extraction negative control (opened prep tube in the PCR hood during sample aliquoting) was included. An additional extraction negative control prep tube remained closed during sample aliquoting. Genomic DNA was extracted as previously described^35^ with the addition of a mechanical lysis step using 0.2 g of 2.8 mm ceramic beads to improve extraction efficiency and without mutanolysin. PCR amplification of the variable 3 and 4 (V3-V4) regions of the 16S rRNA gene was subsequently performed on the extracted DNA from each sample using methods previously described.^36^ In addition to the typical 30 cycles of amplification, each sample was also independently amplified for 40 cycles to confirm any signals detected—these additional amplifications were not used for general analysis as amplification beyond 30 cycles is known to bias sequencing data.^37^ Each reaction contained 5 pmol of primer (341F – CCTACGGGNGGCWGCAG, 806R – GGACTACNVGGGTWTC-TAAT), 200 mM of dNTPs, 1.5μl 50 mM MgCl_2_, 2 μl of 10 mg/ml bovine serum albumin (irradiated with a transilluminator to eliminate contaminating DNA) and 0.25μl Taq polymerase (Life Technologies, Canada) for a total reaction volume of 50 μl. 341F and 806R rRNA gene primers were modified to include adapter sequences specific to the Illumina technology and 6-base pair barcodes were used to allow multiplexing of samples. 16S DNA products of the PCR amplification were subsequently sequenced twice on two separate runs of the Illumina MiSeq platform (2×300bp) at the Farncombe Genomics Facility (McMaster University, Hamilton ON, Canada).

### 16S rRNA gene sequencing and analysis

Primers were trimmed from FASTQ files using Cutadapt^38^ (RRID:SCR_011841) and DADA2^39^ was used to derive amplicon sequence variants (ASVs). Taxonomy was assigned using the Silva 132 reference database.^40^ Non-bacterial ASVs were culled (kingdom Eukaryota, family Mitochondria, order Chloroplast, or no assigned phylum), as was any ASV to which only 1 sequence was assigned.

We performed alpha and beta diversity analyses in R using phyloseq^41^ (RRID:SCR_013080) and tested for whole community differences across groups using vegan’s^42^ (RRID:SCR_011950) implementation of permutational multivariate analysis of variance (PERMANOVA) in the adonis command. These results were visualized via Principal Coordinate Analysis (PCoA) ordination using R’s ggplot2 package (RRID:SCR_014601).^43^ ASV sequences were aligned using DECIPHER^44^ and a GTR+G+I (Generalized time-reversible with Gamma rate variation) maximum likelihood tree phylogenetic tree was constructed with phangorn^45^ using a neighbor-joining tree as a starting point. Significance of alpha diversity was analyzed by linear mixed model with sample type as a fixed effect and participant ID as a random effect. Significance of Bray-Curtis distances between and within sample types was assessed by linear mixed model with sample type comparison as a fixed effect (i.e. neg-neg, neg-fetal, etc.) and participant ID as a random effect.

## Data availability

All sequencing data associated with this study has been made publicly available. 16S rRNA bacterial profiling data generated in this study is available in the NCBI Sequence Read Archive under project ID PRJNA666699.

## Acknowledgements

We thank all the participants that were recruited in this study. We would like to thank Dr. Hanna Brinkmann, Laura Pasura, Laura Maschirow and Alexander Schwickert for assisting with patient recruitments; Mrs. Loreen Ehrlich with sample preparation and Dr. Katharina von Weizsaecker for her external review of the microbiology protocol and advice on protocol improvements. We thank Michelle Shah for performing genomic DNA extractions. K.M.K. is supported by a Farncombe Digestive Health Research Institute Student Fellowship. MGS and DMS are supported by the Canada Research Chairs Program.

## Author Contributions

K.M.K. analyzed sequencing data and wrote the manuscript. M.G. contributed to sample collection and wrote the Study Design and Sample Collection portion of the Methods section. T.A. performed the culture-based analyses. M.M.H. assisted in study design. L.R. assisted in study design and performed V3V4 amplifications and processing of raw sequencing data. M.G.S. assisted in study design and analysis of sequencing data. D.M.S. contributed to data analysis, and manuscript development. T.B. designed the study and contributed to sample collection. All authors discussed the analyses and results and edited the manuscript.

---

**Supplementary Figure 1.**
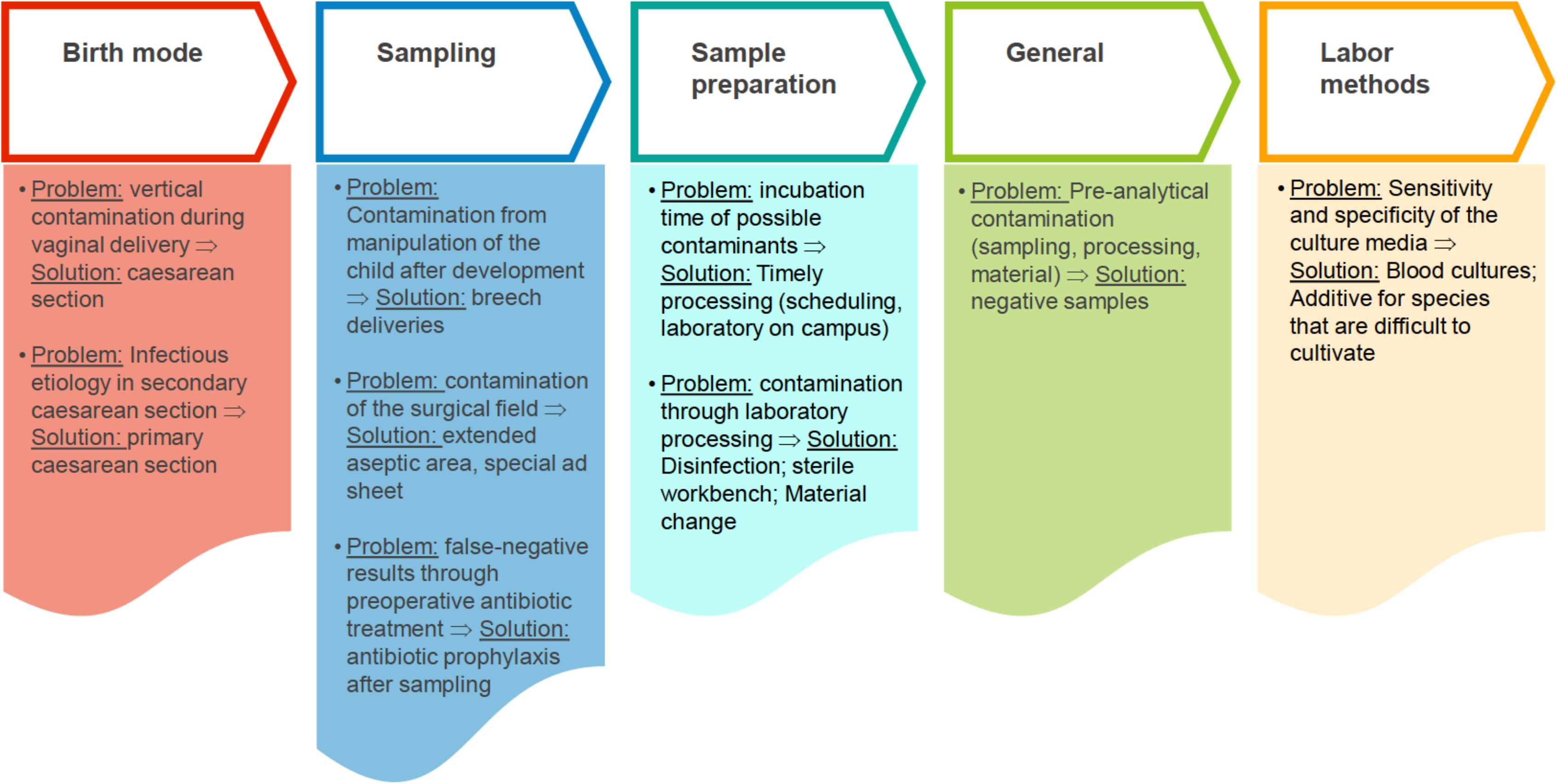
Optimization of collection to reduce contamination.

**Supplementary Figure 2.**
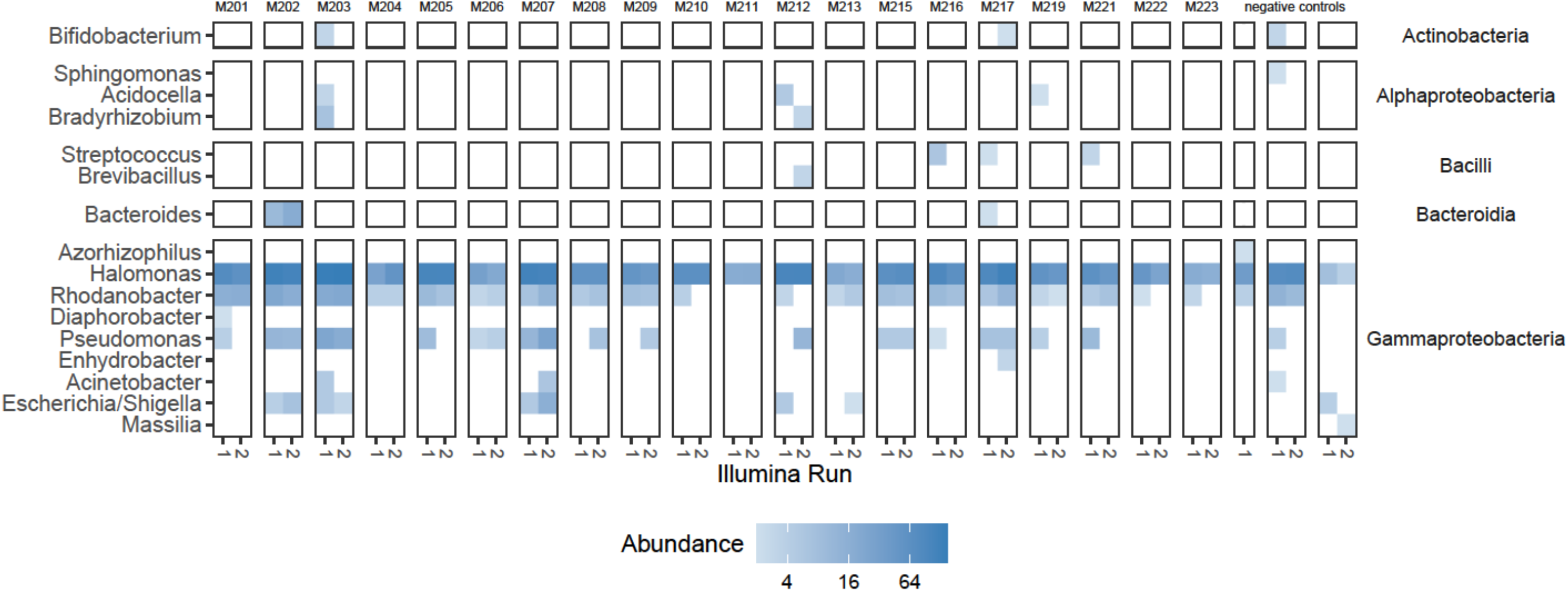
Detection of genera across technical replicates. The abundance (read count) is shown for each sample by sequencing run (run 1 and run 2) for 30 cycles of PCR amplification. Genera are only shown if they were also detected within the same sample’s sequencing data from 40 cycles of PCR amplification.

